# The genital tract microbiota of IVF patients and oocyte donors: a comparative analysis

**DOI:** 10.1101/2024.10.18.619104

**Authors:** Thor Haahr, Peter Humaidan, Andrea Bernabeu, Belen Lledo, Erica L Plummer, Catriona S Bradshaw, Anna Cäcilia Ingham, Jørgen Skov Jensen

**Affiliations:** Department of Clinical Medicine, Aarhus University, Denmark and the Fertility Clinic Skive, Skive Regional Hospital, Denmark; The Fertility Clinic, Instituto Bernabeu, Alicante, Spain; Melbourne Sexual Health Centre, Alfred Health, Melbourne, Australia; School of Translational Medicine, Monash University, Melbourne, Australia; Department of Bioinformatics, Statens Serum Institut, Copenhagen, Denmark; Research Unit for Reproductive Microbiology, Statens Serum Institut, Copenhagen, Denmark

**Keywords:** Vaginal microbiota, Bacterial vaginosis, Cervix, Uterus, Microbiome, IVF, Infertility, oocyte donation

## Abstract

Recent studies have observed an association between genital tract microbiota and reproductive outcomes in patients undergoing in vitro fertilization (IVF). This finding could be caused by ascending dysbiosis from the vagina to the endometrium. The main objective of this study was to compare the microbiota and total bacterial load in the vagina, cervix and endometrium of IVF patients and oocyte donors. Thus, a total of two cohorts were explored: IVF patients (N=27) and healthy oocyte donors (N=26). The microbiota was investigated using 16S rRNA gene sequencing of the V3-V4 hypervariable regions, as well as qPCR for common vaginal species and total bacterial load. The study used a transcervical sampling approach for the endometrial microbiota samples.

The highest bacterial load was seen in the vagina (median 2.6*10^9^; 95% CI 9.7*10^8^ - 7.1*10^9^; 16S rRNA gene copies/swab), followed by the cervix (median 9.0*10^7^; 95% CI 3.3*10^7^ - 2.4*10^8^; 16S rRNA gene copies/swab) and the endometrium (median 8.6*10^6^; 95%CI 3.1*10^6^ - 2.4*10^7^; 16S rRNA gene copies/swab). Only 1% of the observed variation between genital tract samples was explained by patient group (IVF patient or oocyte donor) and less than 1% of the variation between genital tract samples was explained by the anatomical site. Although, findings could be impacted by the transcervical sampling approach, 16S rRNA gene sequencing as well as species-specific qPCR showed a similar microbiota composition present in the cervix and the endometrium when compared to the vaginal microbiota.

## Importance

This study observed no significant differences in the genital tract microbiota composition between IVF patients and oocyte donors who underwent similar ovarian stimulation protocols and were sampled at a fixed time point during midcycle. A bacterial load gradient was observed in the female genital tract with the bacterial load being highest in the vagina, lower in the cervix and lowest in the endometrium. In both qPCR for selected species, and in 16S rRNA gene analyses, we observed a similar microbiota composition when comparing the vagina, cervix and endometrium. Although the endometrial microbiota could be influenced by the transcervical sampling approach, we did not observe “signature” bacterial taxa in the endometrium. Rather, we found that the endometrial microbiota was a reflection of the vaginal microbiota.

## Introduction

Historically, the uterine cavity of healthy women has been perceived as sterile. However, an endometrial microbiota has been described, using both culture(1, 2) and culture independent methods(3–6). Some studies reported that the endometrial microbiota resembles that of the cervico- vaginal microbiota(1–3, 5, 6), whereas others found a unique microbiota with ‘signature taxa’ in the endometrium and upper genital tract(4, 7, 8). Importantly, many of the unique ‘signature taxa’ reported in endometrial microbiota studies have also been observed in negative/blank controls(9, 10), highlighting that the endometrial microbiota is a low biomass environment in which stringent experimental controls are needed to distinguish genuine biological signals from contaminants(11, 12).

Interestingly, recent studies have correlated genital tract dysbiosis with poor reproductive outcomes in IVF patients(5, 13–16), and it could be hypothesized that the underlying aetiology is an ascending infection from the vagina to the endometrium. In support of this, Swidsinski *et al*. reported an increased risk of a *Gardnerella* dominated biofilm attached to the endometrium of women with bacterial vaginosis (BV)(17). Additionally, a correlation between endometrial dysbiosis and chronic endometritis has also been reported(18).

A prior retrospective case-control study used 16S rRNA gene sequencing to compare the vaginal, cervical and endometrial microbiome of 15 women with a history of infertility with that of 16 fertile controls (6). The study reported that in general the samples clustered on the level of the individual rather than on the anatomical site. However, the study was limited by its retrospective design and small sample size. Thus, the aims of the present study were to investigate the microbiota of the vagina, cervix and endometrium in IVF patients and healthy oocyte donors using qPCR and 16S rRNA gene sequencing to: i) determine differences in microbiota composition between IVF patients and oocyte donors and ii) quantify bacterial composition and load at the three anatomical sites with a particular focus on women with abnormal vaginal microbiota (AVM). As described previously, AVM is highly correlated with BV (diagnosed by Nugent Score) and is defined using qPCR as a vaginal microbiota with high bacterial loads of *Gardnerella* (≥5.7*10^7^ 16S rRNA gene copies per mL) and/or *Fannyhessea (F.) vaginae* (≥5.7*10^6^ 16S rRNA gene copies per mL)(13).

## Results

In total, 27 IVF patients and 26 oocyte donors were recruited in two independent prospectively registered cohorts. Baseline characteristics for the IVF patients and the oocyte donors are shown in **Table 1**. Vaginal, cervical and endometrial samples were taken and a total of 157/159 samples were analysed, as two vaginal swabs (Study ID, IVF3 and IVF18) were lost. Study ID IVF14 reported the use of a *L. gasseri* containing vaginal probiotic and had *L. gasseri* detected in all anatomical sites, whereas the other patient reporting use of vaginal probiotics had AVM and did not specify the specific product used (Study ID, IVF26).

**Table 1:**
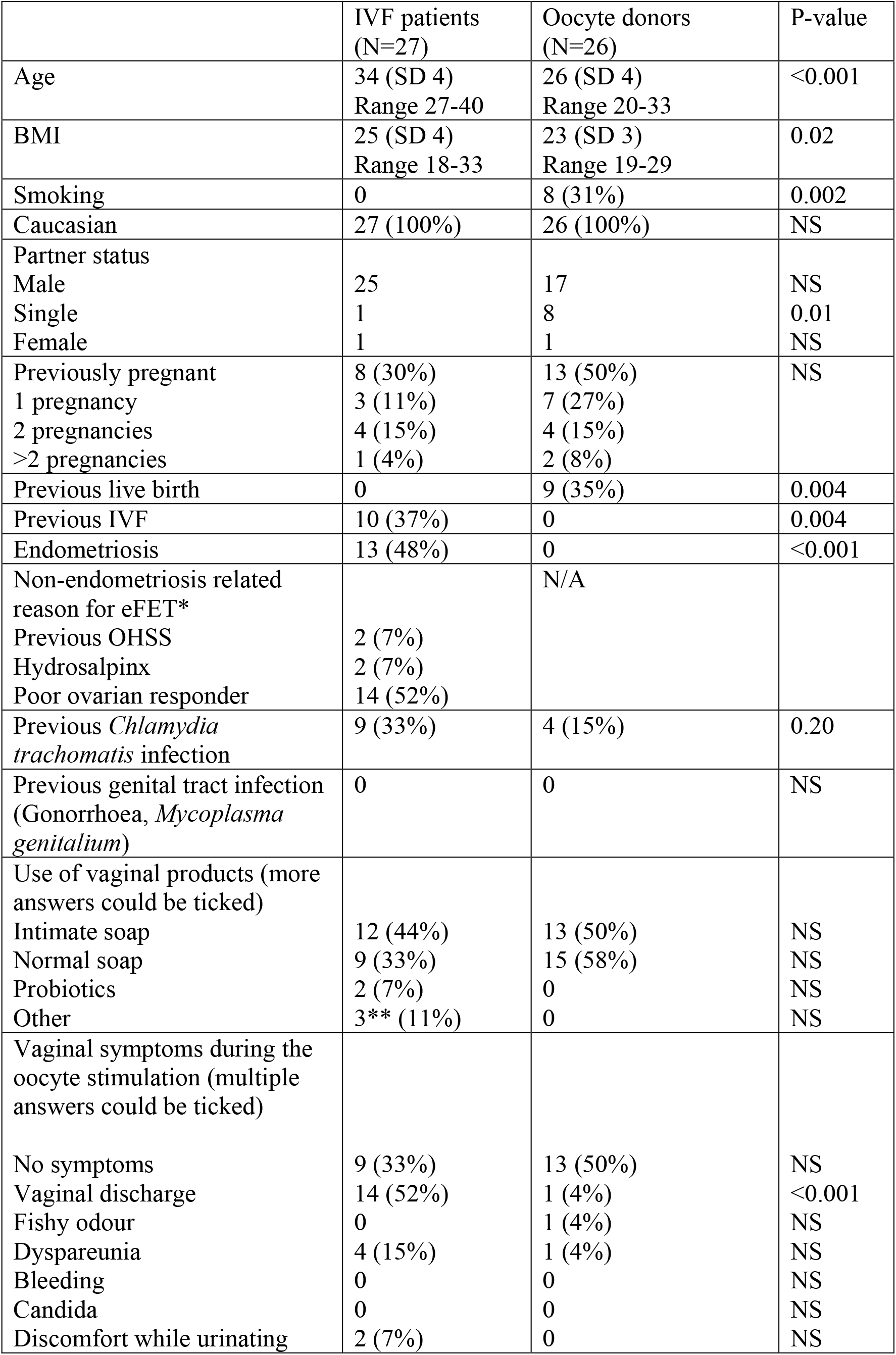
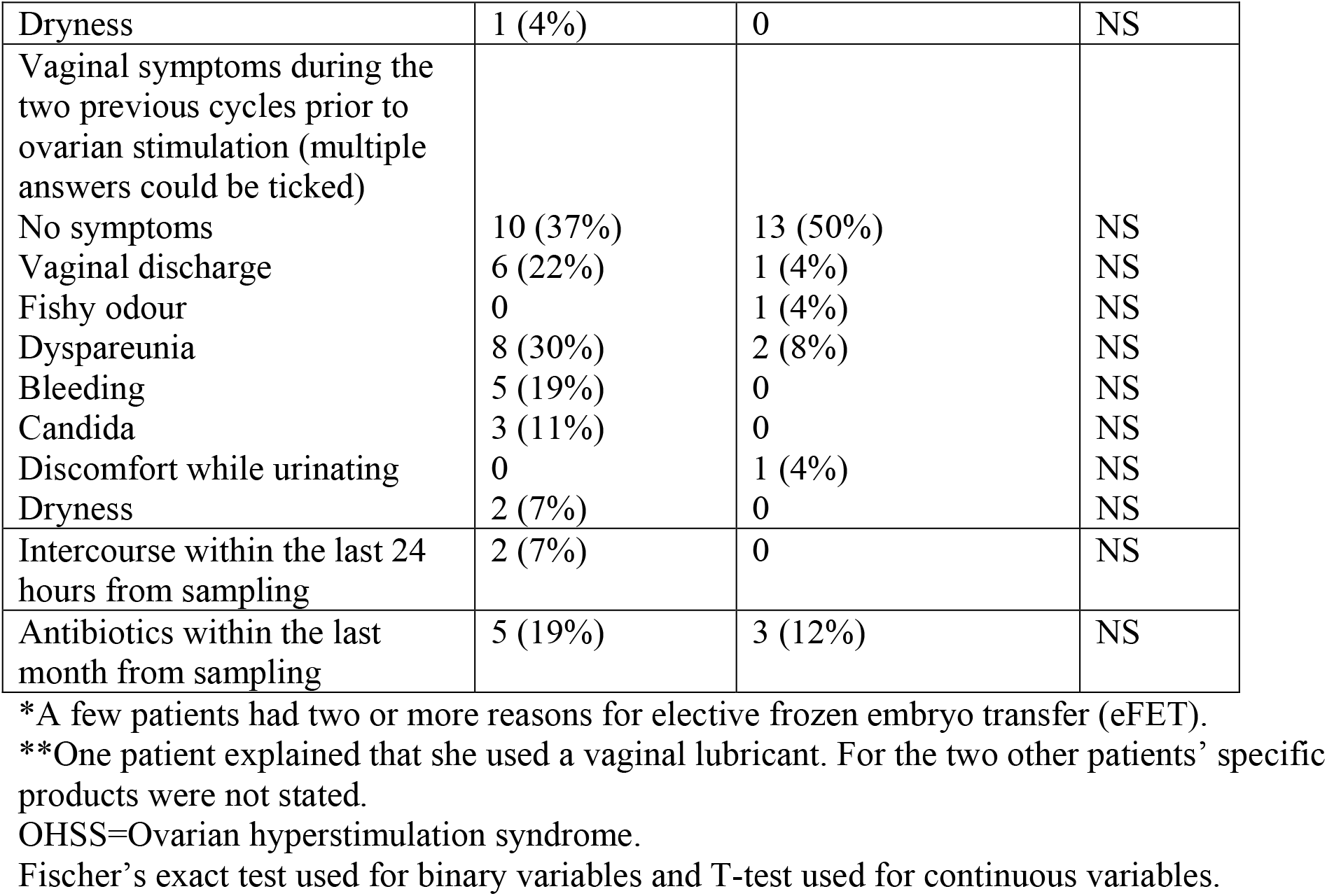
Baseline characteristics of the two cohorts.

### Comparison between IVF patients and oocyte donors

In **Figure 1**, the genital tract microbiota was compared between oocyte donors and IVF patients (who had suffered from at least one year of non-male-factor infertility (N=22)) by principal component analysis (PCA). Samples clustered mainly on the individual level of the woman, explaining 84% (Jaccard) - 89% (Bray-Curtis) of the variation (adonis, P<0.001), whereas being either an oocyte donor or IVF patient explained only 1% of the variation with both Jaccard and Bray-Curtis index (adonis, P=1.000). The overall prevalence of AVM in the study population was 38% (20/53) and was not significantly different between IVF patients 9/27 (33%) and oocyte donors 11/26 (42%), P=0.50. Moreover, there were no statistically significant differences between IVF patients and oocyte donors in species-specific qPCR bacterial loads, including common *Lactobacillus spp.*, *Gardnerella* and *F. vaginae* (data not shown).

**Figure 1:**
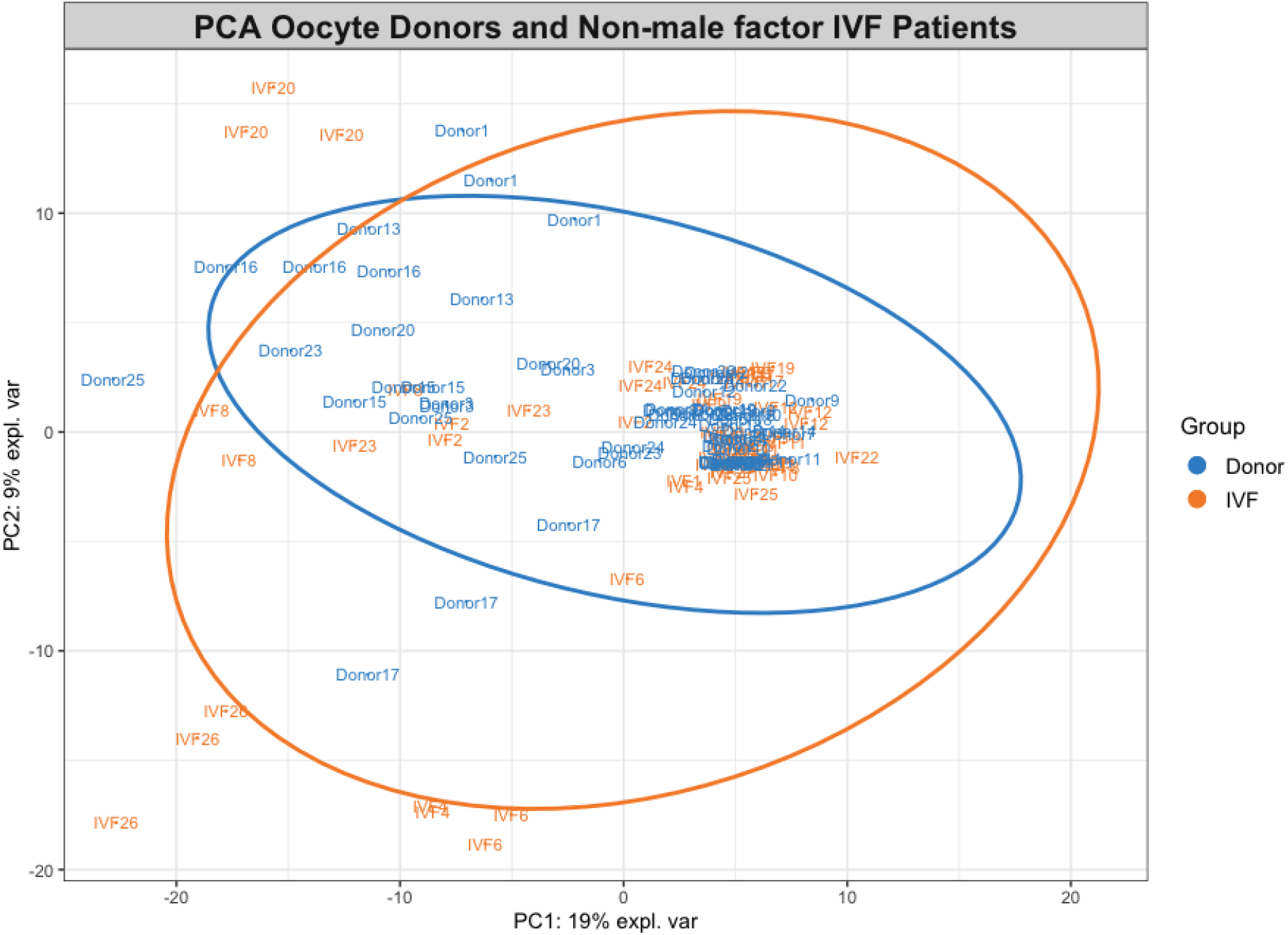
Principal component analysis comparing oocyte donors and non-male factor IVF patients. Figure 1 legend: Overlapping 95% CI ellipses between samples from oocyte donors (N=26) and female IVF patients suffering from at least one year of non-male factor infertility (N=22). Samples from the three sample sites cluster on the level of the individual as can be seen with the labelled study ID. Missing samples were due to either low read counts (<218 reads, N=15) or missing samples (N=2)

### qPCR results of bacterial ascension from the vagina to the endometrium

The paired vaginal, cervical and endometrial samples among oocyte donors were analysed for bacterial load by a general probe for 16S rRNA gene copies. In both the mixed statistical model and using non-parametric statistics, there was a statistically significant gradient in total bacterial load comparing the individual sample sites. The highest total bacterial load was observed in the vagina, followed by the cervix and the lowest bacterial load was found in the endometrium, **table 2 and Figure S1**. In the vagina, the total bacterial load in non-AVM samples was significantly lower compared to AVM samples for oocyte donors, **Figure S2** (P<0.001). Likewise, in the cervix, the bacterial load was lower for oocyte donors without AVM compared to oocyte donors with AVM (P=0.006). There was no statistically significant difference in endometrial bacterial load between oocyte donors with AVM and oocyte donors without AVM (P=0.16).

**Table 2:** Comparison of the qPCR bacterial loads by the statistical models described in methods.

|  | **Vagina**<br>**(Median, 95%**<br>**CI)** | **Cervix**<br>**(Median, 95%**<br>**CI)** | **Endometrium**<br>**(Median, 95%**<br>**CI)** | **V/C**<br>**Ratio** | **V/E**<br>**Ratio** | **C/E**<br>**Ratio** |
| --- | --- | --- | --- | --- | --- | --- |
| **Total 16S**<br>**bacterial load**<br>**(N=26)** | **2.6\*10<sup>9</sup>**<br>(9.7*10<sup>8</sup> -<br>7.1*10<sup>9</sup>) | **9.0\*10<sup>7</sup>**<br>(3.3*10<sup>7</sup> -<br>2.4*10<sup>8</sup>) | **8.6\*10<sup>6</sup>**<br>(3.1*10<sup>6</sup> -<br>2.4*10<sup>7</sup>) | **29** | **305** | **10** |
| ***L. crispatus*, *L.***<br>***jensenii*, *L.***<br>***iners* and *L.***<br>***gasseri***<br>**(N=51)** | **2.7\*10<sup>7</sup>**<br>(1.2*10<sup>7</sup> -<br>6.1*10<sup>7</sup>) | **3.0\*10<sup>5</sup>**<br>(1.3*10<sup>5</sup> -<br>6.8*10<sup>5</sup>) | **5.3\*10<sup>4</sup>**<br>(2.3*10<sup>4</sup>-1.2*10<sup>5</sup>) | **90** | **509** | **6** |
| ***Gardnerella***<br>**and *F. vaginae***<br>**(N=51)** | **5.4\*10<sup>6</sup>**<br>(1.1*10<sup>6</sup> -<br>2.5*10<sup>7</sup>) | **1.4\*10<sup>5</sup>**<br>(2.9*10<sup>4</sup>-<br>6.7*10<sup>5</sup>) | **6.6\*10<sup>4</sup>**<br>(1.4*10<sup>4</sup>-3.2*10<sup>5</sup>) | **39** | **82** | **2**<br>(P=0.09) |
| **All BV**<br>**associated**<br>**bacteria\***<br>**(N=53)** | **8.3\*10<sup>6</sup>**<br>(2.1*10<sup>6</sup> -3.2*10<sup>7</sup>) | **2.1\*10<sup>5</sup>**<br>(5.3*10<sup>4</sup>-8.0*10<sup>5</sup>) | **1.4\*10<sup>5</sup>**<br>(3.5*10<sup>4</sup>-5.4*10<sup>5</sup>) | **40** | **60** | **1.5**<br>(P=0.32) |

\*BVAB1, BVAB2, BVAB3, *Prevotella*, *Sneathia*, *Leptotrichia*, *F.vaginae*, *Gardnerella*, *M.hominis*

The species-specific qPCR results were available for all 53 women. The vaginal, cervical and endometrial samples are visualized in **Figure 2**. There was no statistical difference in a combined group of species-specific BV associated bacteria, comparing the cervical and endometrial samples; however, all other sample site comparisons corroborated the aforementioned statistically significant bacterial gradient between sample sites for both the group of BV associated bacteria and for the group of lactobacilli combined, **table 2**. The combination of groups was performed to make parametric statistics possible and to avoid the many 0 copies/swab for the individual species- specific qPCR targets. Importantly, the negative controls had 0 gene copies/swab for all the species- specific qPCR targets, yielding confidence in the low qPCR loads generally reported in both cervical and endometrial samples.

**Figure 2.**
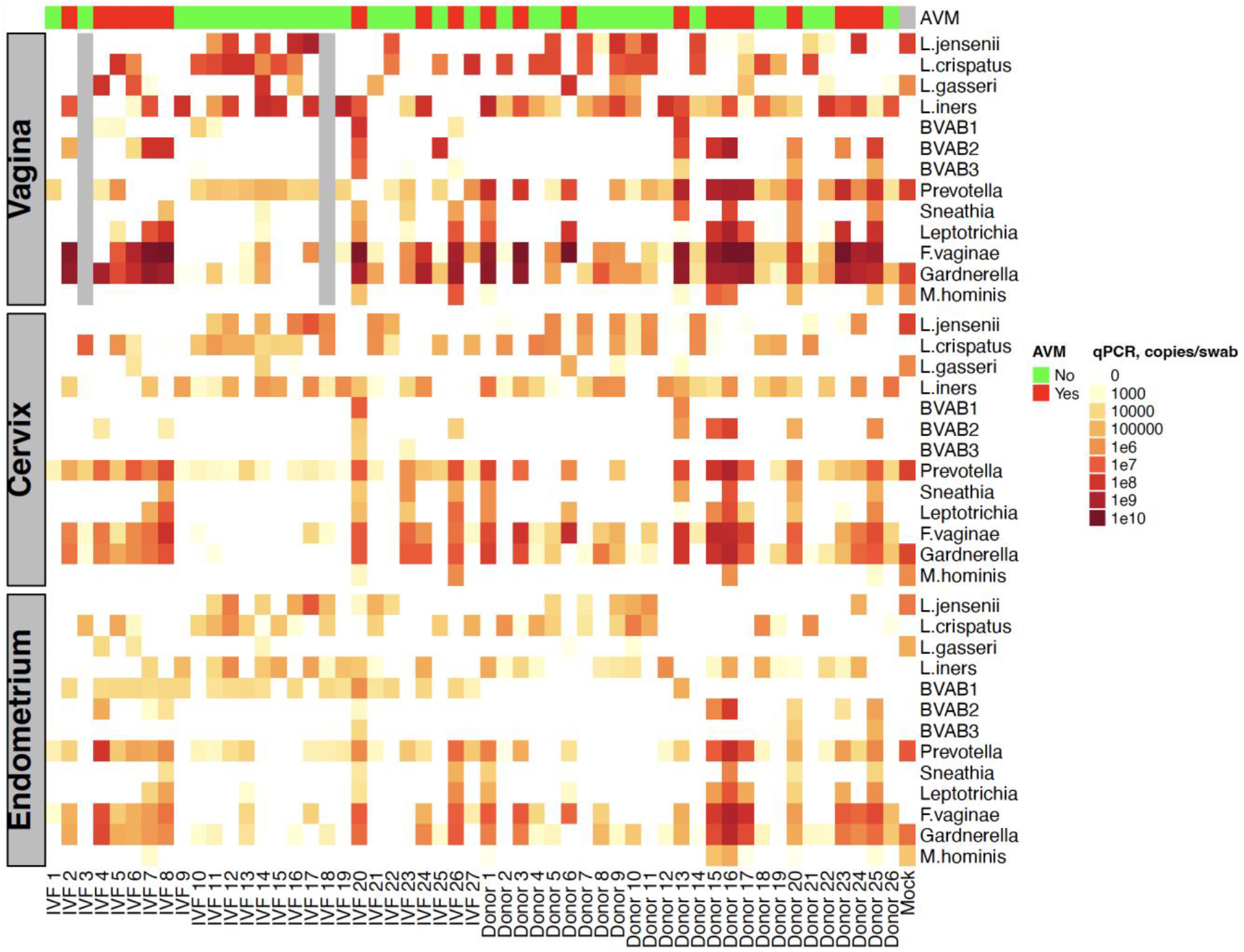
qPCR bacterial load of selected species in vaginal, cervical, endometrial and mock samples. Figure 2 legend: Vaginal, cervical and endometrial gene copies per swab in qPCR analysis. Aligned vertically in a heatmap for each study participant. Gray bars represent two missing vaginal samples for IVF3 and IVF18. Mock is a vaginal mock community that was processed with the respective samples as described in the supplement. AVM=abnormal vaginal microbiota determined by qPCR.

In women with AVM, there was a statistically lower *Gardnerella* load in the endometrium (median 3.7*10^6^ gene copies/sample; Interquartile range (IQR) 3.7*10^5^-3.8*10^6^) compared to the cervix (median 1.2*10^7^ gene copies/sample; IQR 5.4*10^6^-3.8*10^7^) (P=0.015). However, the *F. vaginae* load was not statistically different when comparing cervical and endometrial samples, (P=0.33). In general, patients without AVM had few or no copies of *Gardnerella* (median 0; range 0-1.2*10^8^ gene copies/sample) and *F. vaginae* (median 0; range 0-8.1*10^6^ gene copies/sample) in the cervix and the endometrium (*Gardnerella*; median 441; range 0-8.9*10^4^ gene copies/sample and *F. vaginae*; median 0; range 0-7.8*10^4^ gene copies/sample). However, as visualized in **Figure S3** some individual patients had a higher *Gardnerella* and *F. vaginae* load in the endometrium when compared to the cervix (e.g., participants with study ID: IVF4, D17 and D23).

When stratified by individual sample site, we observed significantly higher loads of *L. crispatus* and *L. jensenii*, but not *L. iners* and *L. gasseri,* in non-AVM compared to AVM samples, **Figure S4**. Moreover, there was a significantly lower number of *L. crispatus* gene copies in the endometrium of women with AVM (median 81; IQR 0-110 gene copies/per swab) compared to women without AVM (median 1.7*10^4^; IQR 110-9.7*10^4^ gene copies/per swab), P=0.001.

### The genital tract microbiome by 16S rRNA gene sequencing

There was a clear correlation between the total 16S rRNA gene copy/swab number in qPCR and the samples excluded due to a low number of sequence reads, Figure S1. Heatmaps were constructed to visualize the similarity between vaginal, cervical, and endometrial microbiota in IVF patients (**Figure 3**) and oocyte donors (**Figure 4**). As visualized in the PCA plot, **Figure 5**, only 0.6-0.7% (Jaccard and Bray-Curtis in adonis function, respectively) of the variation was explained by anatomical site. Since comparing bacterial composition between patients by adonis comes with limited statistical power due to a high number of groups with only three observations each (i.e., three sample sites), we additionally compared the ordination plots, using procrustes analysis. Procrustes analysis confirmed a highly significant correlation between anatomical sites, Sum of Squares =0.29, P=0.001 for vaginal to cervical ordination and Sum of Squares=0.49, P=0.001 for vaginal to endometrial ordination.

**Figure 3:**
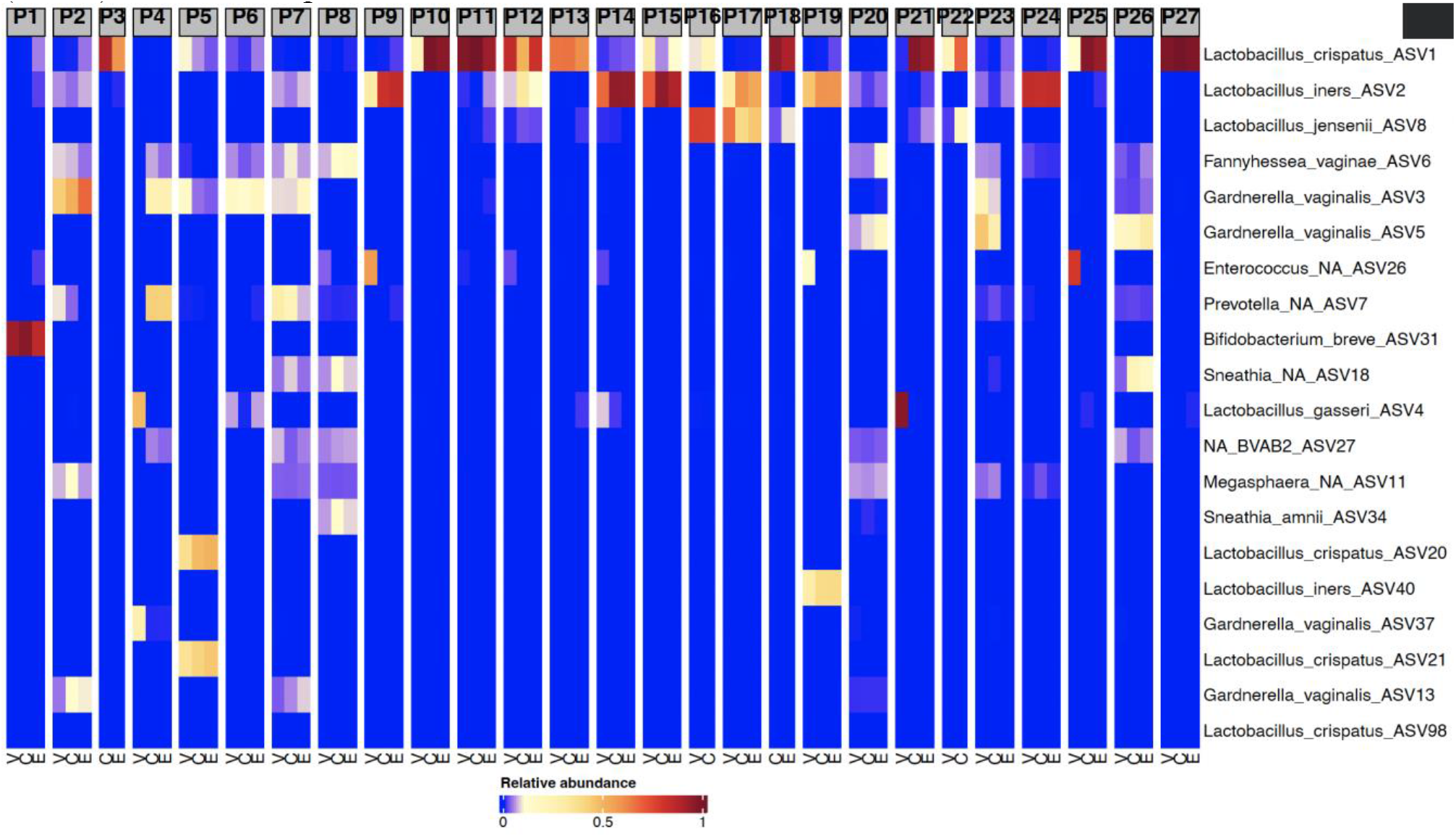
Individual heatmaps of all IVF patients showing top 20 amplicon sequence variants (ASVs) from all sample sites as based on relative abundance. Figure 3 legend: Heatmaps subset to individual level to visualise microbiota composition per sample site. P1=IVF patient 1 etc. V=Vagina, C=Cervix and E=Endometrium. Missing samples were due to either low read counts (<218 reads) or missing samples (N=2).

**Figure 4:**
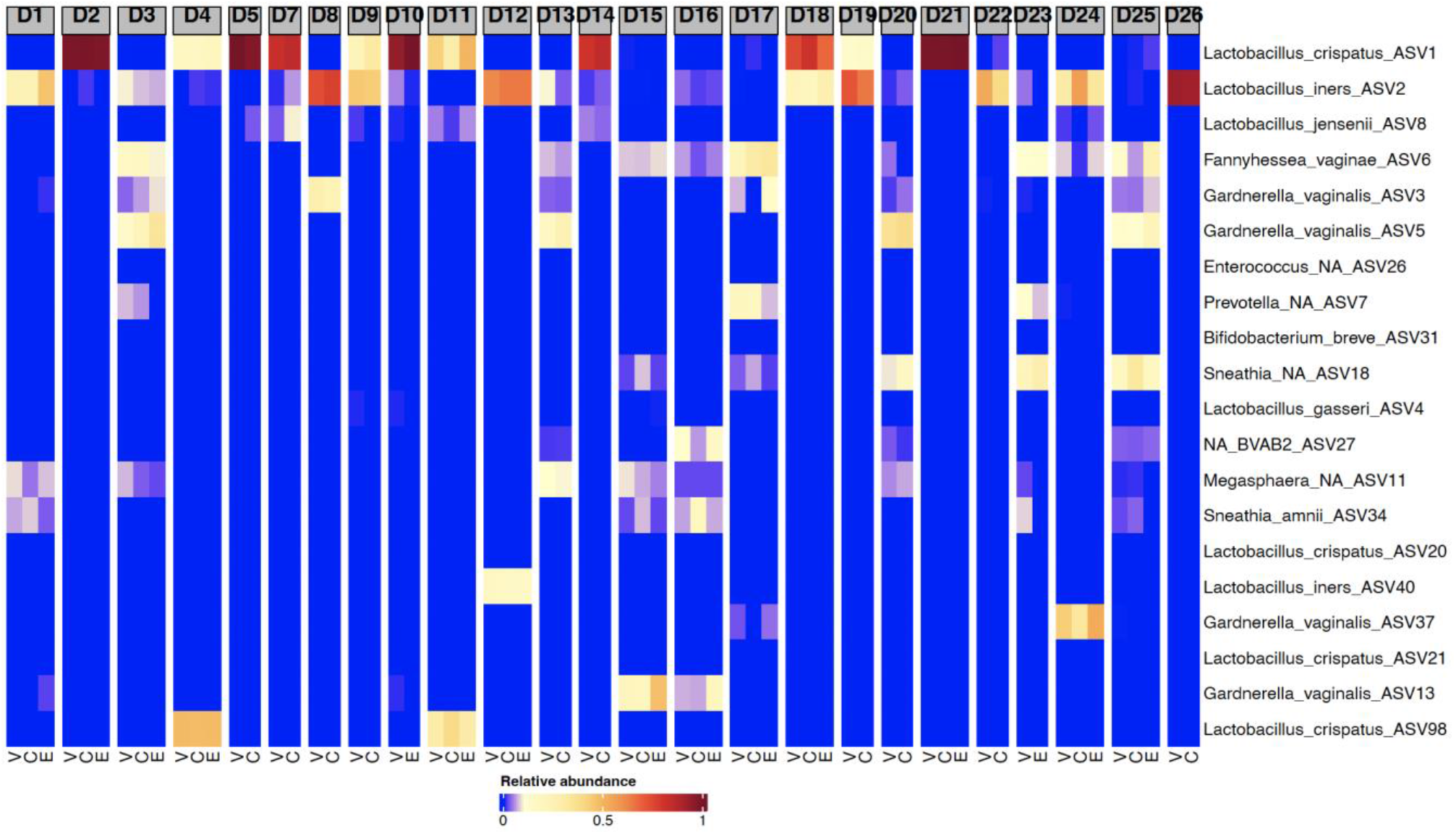
Individual heatmaps of all oocyte donors showing top 20 amplicon sequence variants (ASVs) from all sample sites as based on relative abundance. Figure 4 legend: Heatmaps subset to individual level to visualise microbiota composition per sample site. D1=Donor 1 etc. V=Vagina, C=Cervix and E=Endometrium. Missing samples were due to low read counts (<218 reads). For Study ID D6 only the vaginal site had >218 reads so that participant was omitted from this figure.

**Figure 5:**
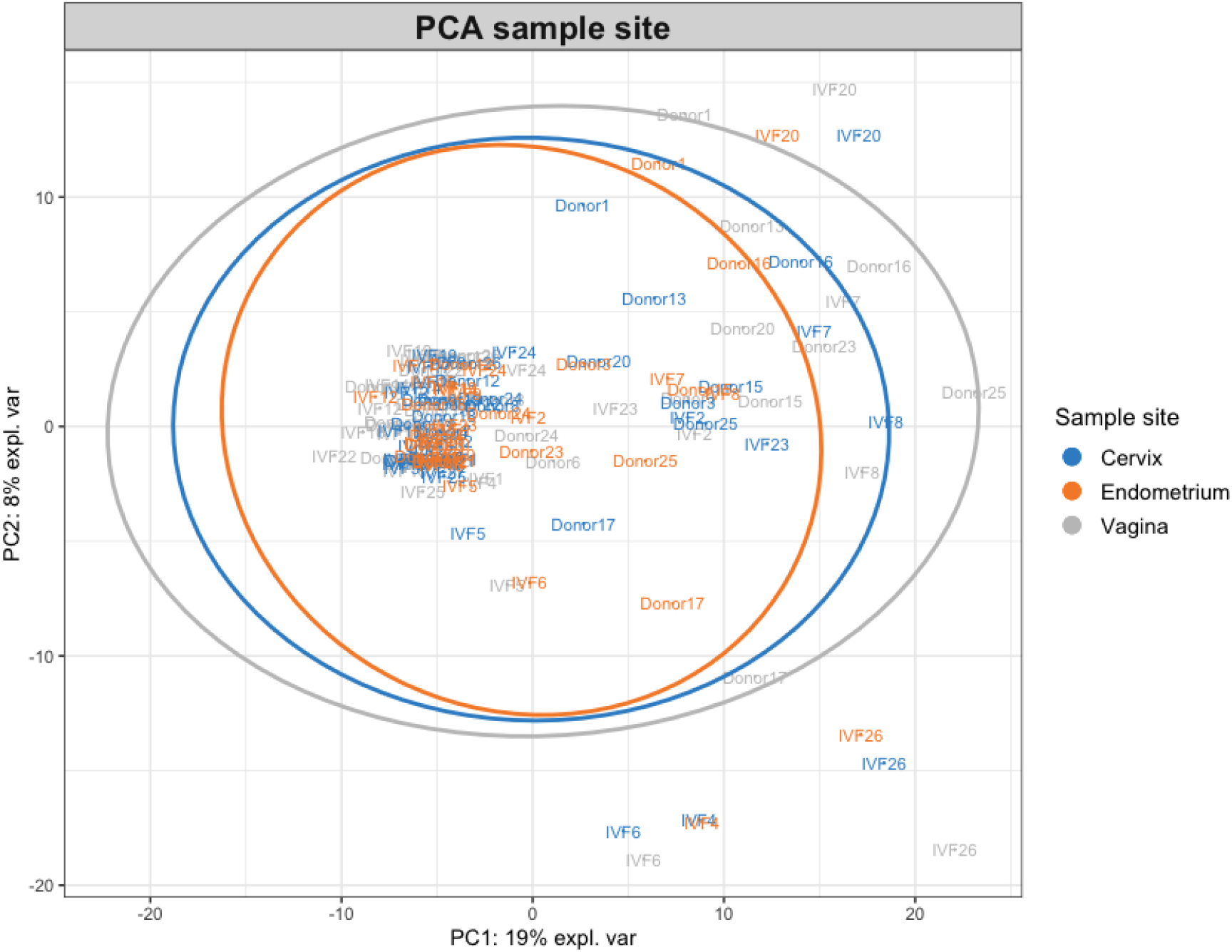
Principal component analysis on vaginal, cervical and endometrial samples from all IVF patients and oocyte donors. Figure 5 legend. Overlapping 95% CI ellipses between sample sites. Samples cluster on the level of the individual as can be seen with the labelled study ID. Missing samples were due to either low read counts (<218 reads, N=15) or missing samples (N=2)

## Discussion

### Main findings

We used a transcervical sampling approach to compare the genital tract microbiota in IVF patients and oocyte donors at the time of oocyte retrieval. The main findings were: i) the genital tract microbiota of IVF patients was not statistically different from that of oocyte donors; ii) the individual variability of the genital tract microbiota between women was greater than the variability between the sample sites investigated; iii) a bacterial gradient of similar composition was present ranging from high bacterial loads in the vagina, to less in the cervix and least in the endometrium; and iv) women with AVM had a significantly lower *L. crispatus* load in the endometrium.

### Interpretation

In general, there is a scarcity of publications investigating the composition of the endometrial microbiota. Thus, the evidence for a genuine endometrial microbiota in reproductive age women can be divided into studies investigating transcervical samples, which cannot completely exclude contamination, and studies investigating endometrial samples from hysterectomised women, which could be considered more or less contamination free(3, 11, 19). However, studies of endometrial samples from hysterectomized women may not resemble the endometrial microbiota of e.g. infertile patients.

One previous study also used the Tao Brush™ sampling device to investigate the endometrial microbiota(7). In that study, the V1-V2 regions of the 16S rRNA gene was sequenced, and the authors reported high relative abundances of either *L. crispatus* or *L. iners* in 7/19 endometrial samples in women suffering from recurrent pregnancy loss or recurrent implantation failure. However, the study did not observe a high relative abundance of *Gardnerella* in any sample. This is likely explained by the use of primers targeting the V1-V2 regions which have a known bias against detecting *Gardnerella*(20), compared to the V3-V4 regions used in our study.

Winters *et al*. (2019) collected endometrial swab samples from women (median age 45, IQR 41- 49.5) undergoing abdominal hysterectomy for fibroma(11), and reported that the endometrial bacterial biomass from 40% (10/25) of women was indistinguishable from the bacterial biomass in negative controls. However, 60% (15/25) of women had an endometrial bacterial biomass that exceeded negative controls and was similar to the cervical bacterial biomass. Moreover, authors concluded that *Lactobacillus spp.* were rare in the endometrium despite a high abundance in the vagina as measured by both 16S rRNA gene sequencing and species-specific qPCR. Unfortunately, that study did not report the diagnosis of BV in which endometrial bacteria might be more likely to be present. In fact, Swidsinski *et al*. found increased odds of endometrial infection in women with BV diagnosed by a FISH method compared to women without BV by the same method, Odds Ratio (OR) 5.7 (95% CI, 1.8–18.3, P = 0.002)(17). Additionally, the Winters *et al*. study(11) found that the cervical microbiota around the external cervical os is a low biomass microbiota.

In the present study, we partly corroborated these results, reporting that the microbial biomass from the cervix is lower compared to the vagina for total bacterial load and most individual bacterial targets in the qPCR analyses. An interesting finding is that the lactobacilli gradient from the vagina to the endometrium is more pronounced compared to the gradient for the BV associated bacteria. This observation could be speculated to have its origin in lactobacilli not being capable of populating the endometrium when compared to the BV associated bacteria, which might have a survival advantage at the higher uterine pH around 7(21).

Moreno and colleagues (2016) investigated the microbiota of endometrial fluid, extracted via a transcervical catheter in women undergoing IVF and embryo transfer after endometrial receptivity testing(5). By using a cut-off level of 90% relative abundance of *Lactobacillus* reads for stratifying IVF patients being *Lactobacillus* dominant or not, the authors reported a 59% (10/17) and 7% (1/15) live birth rate per embryo transfer, respectively. This intriguing finding was corroborated by the same research group in 2022, using 16S rRNA gene sequencing of endometrial samples. The authors reported that *Atopobium* (now *Fannyhessea*), *Gardnerella*, *Staphylococcus* and *Streptococcus* genera were associated with a poor live birth rate in IVF patients whereas a *Lactobacillus* dominant endometrial microbiota was associated with significantly higher live birth rates(19). In contrast, Hashimoto et al. (2019) reported a 38% (26/68) ongoing pregnancy rate per blastocyst transfer in IVF patients with above 80% relative abundance of *Lactobacillus spp.* and *Bifidobacterium spp.* in the endometrium as compared to a 45% (14/31) ongoing pregnancy rate in IVF patients not dominated by *Lactobacillus spp.* and *Bifidobacterium spp.* (22). Thus, results are conflicting regarding the role of the endometrial microbiota in relation to IVF outcomes.

### Strengths and limitations

A previous study compared upper genital tract microbiota of infertile patients to fertile controls(6). Compared to that study which recruited historical infertile patients(6), the strength of our study is the inclusion of IVF patients experiencing at least one year of infertility and who underwent an elective frozen embryo transfer cycle. It is also a strength that all participants underwent similar ovarian stimulation protocols prior to sampling at a fixed time during the cycle in which the cervical canal was fully dilated. Although restricted to a Caucasian origin, it is a limitation that the IVF patients and oocyte donors were from Denmark and Spain, respectively. Moreover, the fertility of oocyte donors in this study was not always evident with regards to a previous live birth, albeit oocyte donors would generally be considered to have a good fertility potential, based on a high number of antral follicles and the absence of endometriosis, intrauterine malformations and other known reproductive disorders.

Importantly, we used stringent experimental controls at all steps and included qPCR analyses, which are optimal for quantification of the bacterial load of specific bacterial species(23). A limitation of this study, however, was that we did not have enough material for the qPCR 16S total abundance measurement in the samples from the IVF cohort. Thus, sample size was limited for this comparison. Further, our inclusion of experimental controls enabled us to be confident in the resulting microbiome composition of low biomass endometrial and cervical samples. The cervical swab used in the present study also sampled parts of the ectocervix and should be interpreted as the microbiota of that anatomic location *in situ*. The purpose of the L-shaped flocked swab was to remove the cervical mucus around the external os and in the distal 1 cm of the endocervix, which minimized the amount of cervical microbiota that could be transferred into the uterus by the transcervical endometrial sampling device, TAO brush™. However, with transcervical endometrial samples, it is not possible to completely avoid pushing a small amount of endocervical mucus/microbiota into the endometrium during sampling. The blank tip of the TAO brush™ may have carried a small amount of cervical mucus which could have impacted the endometrial microbiota analysis of the present study. However, it is unlikely that this small amount of cervical mucus influenced results substantially. As an example, some women had even more abundant *Gardnerella* and *F. vaginae* gene copies in the endometrial samples compared to the flocked cervical swab as seen by both qPCR and 16S rRNA gene sequence analysis. In this aspect, it is considered that the endometrial samples of the present study were large and blood-filled tissue samples which would dilute the small amount of cervical microbiota from the tip of the TAO brush™.

Taken together, the similarity in bacterial composition between endometrial, vaginal and cervical samples observed in the present study could be due to i) physiological ascension of bacteria from the vagina to the endometrium – especially at the time of ovulation where the cervix is fully dilated, ii) increased instrumentation during ovarian stimulation, and/or iii) an effect of minute amounts of endocervical bacteria, contaminating the endometrial sample. However, as discussed above we consider that the relatively large endometrial tissue sample obtained in the present study would dilute the relatively small amount of cervical microbiota. Therefore, the high loads of BV associated bacteria seen – sometimes more than at the level of the cervix compared to the endometrium - could be considered as evidence for a low abundant endometrial microbiota during the midcycle in some, but potentially not all, reproductive age women undergoing ovarian stimulation.

## Methods

### Eligibility criteria, recruitment, and ovarian stimulation

The present study consisted of two cohorts, initially compared in a case-control design, followed by a cross-sectional evaluation of the genital tract microbiota. In the first cohort, we invited women undergoing oocyte retrieval before an elective frozen embryo transfer cycle in a public IVF centre in Denmark. The cohort consisted of patients with endometriosis (N=13), defined as ultrasound visible endometriosis cysts and/or previous surgery for endometriosis and/or a typical endometriosis anamnesis. Previous ovarian hyperstimulation (OHSS) patients (N=2), patients with hydrosalpinx (N=2, and poor ovarian responder patients (N=14) undergoing elective double ovarian stimulation (Duostim) for embryo accumulation were also included(24, 25). A total 22/27 (81%) of IVF patients had experienced at least one year of non-male factor infertility. The remaining five IVF patients had male factor, or no male partner and their samples were excluded from the principal component analysis, comparing IVF patients and oocyte donors (Figure 1). The exclusion criteria for the IVF cohort included presence of intrauterine malformations, HIV and/or Hepatitis B and C, HPV, CIN 2 or higher, *Chlamydia trachomatis* infection or BMI>35. Patients were also excluded if they had uncontrolled concomitant diseases, e.g., diabetes or hypertension. In endometriosis patients, ovarian stimulation was performed in a GnRH antagonist protocol starting with an FSH dose between 150-300IU according to age, antral follicle count and BMI. The FSH dose was adjusted according to ovarian response on cycle day 6-7. Patients undergoing Duostim cycles were stimulated with corifollitropin alpha in a fixed protocol as previously reported(25). All patients underwent ovulation trigger with GnRH agonist 36h prior to sampling.

The second cohort consisted of healthy young women, undergoing ovarian stimulation for oocyte donation at the Instituto Bernabeu, Alicante, Spain. The inclusion criterion for oocyte donors was a regular menstrual cycle. Exclusion criteria included use of antibiotics within 1-month, use of vaginal products other than for menstrual hygiene within 1-month, uterine malformations, HIV, Hepatitis B or C positivity, HPV CIN 2 or higher and *C. trachomatis* infection. All oocyte donors were co-treated with a GnRH antagonist during stimulation with exogenous FSH, dosed according to age, antral follicle count and BMI. Ovulation trigger was performed with a bolus of GnRH agonist 36h prior to sampling. Baseline demographic data were captured, using a questionnaire in REDCap™(26, 27) at the respective clinics, hosted at a secure server at Aarhus University, Denmark. All participants provided written informed consent prior to inclusion.

### Sampling procedure

All samples were collected at the time of oocyte retrieval (OR). As part of the standard treatment, patients were instructed not to have unprotected intercourse within 36h prior to OR/sampling. A flocked vaginal swab (Copan, Brescia, Italy) was collected from the posterior fornix during speculum examination prior to normal OR procedures and placed in ESwab® transport medium (Copan, Product number: 480CE). Subsequently, a sterile L-shaped cervical swab (Copan, Product number: 606CS01L) was inserted 1 cm into the cervical canal and rotated after which it was placed in eNat® transport medium. Following this, local anaesthetics was applied in the vagina as part of the normal transvaginal OR procedure.

Before the transvaginal OR, an endometrial sample was obtained, using a sterile TAO brush™ (COOK® Medical) protected by a sheath placed in the cervix. The sheath protects the brush while passing through the cervix. Once in the uterus, the brush was advanced from the sheath to the endometrium in order to collect the endometrial sample with a 360-degree circular movement of the brush. The TAO brush™ collects a 6mm diameter and 3.5cm long biopsy from the endometrial tissue, i.e., not only a “surface brush”. Finally, the brush was retracted and protected by the sheath while passing back through the cervix. The brush was removed from the sheath and cut off into an ESwab® tube. OR was performed according to standard procedures, including antibiotic prophylaxis for selected patients and oocyte donors after sample collection. Sterile gloves were used during all steps and utmost care was exercised not to contaminate samples by touching non-sample site places.

Vaginal swabs, cervical swabs and endometrial biopsies were immediately frozen at −80°C until analysis, with the exception that, vaginal samples from IVF patients were sent at ambient temperature to a central laboratory for immediate AVM diagnosis as described previously(13). Hereafter vaginal samples from IVF patients were stored at −80 until further microbiota analyses.

### DNA Purification, 16S rRNA gene sequencing and Bioinformatics pipeline

DNA extraction was performed according to a standardized method described in supplement. Negative controls for H_2_O, tissue lysis buffer, bacterial lysis buffer, PBS, Tao brush rotated in the air in the oocyte retrieval room, eNat® buffer (Copan) and ESwab® buffer (Copan) were extracted and analysed on the same plates as the relevant samples. The quality of the DNA extraction process was documented by simultaneous extraction of a vaginal mock community (ATCC® MSA-2007™, LGC Standards, Teddington, UK). Primers and PCR conditions prior to sequencing on the Illumina Miseq are provided in detail in the supplement. Denoising and Amplicon Sequence Variant (ASV) inference was performed using DADA2 (v1.12.1 in R v3.5.1). Potential contaminants were identified and removed as stated in the supplementary material. Based on rarefaction curves and the mean number of sequence reads in the 21 negative controls, we excluded all samples with less than 218 reads. This resulted in the exclusion of 15 samples (12 endometrial and three cervical samples). More details of the bioinformatic pipeline can be found in the supplement.

### qPCR

PCR-methods were used to quantitatively detect Gardnerella, Fannyhessea (F.) vaginae, L. iners, L. crispatus, L .jensenii, L. gasseri, BVAB 1, BVAB 2, M. indolicus, Prevotella spp., Sneathia sanguinegens, Sneathia amnii, and Mycoplasma hominis, as described previously(28–30). Recently, Gardnerella vaginalis has been divided into different Gardnerella spp.(31). In this aspect, the qPCR assay used for Gardnerella in the present study targets all the Gardnerella spp. described, but does not distinguish them(31). Thus, the term Gardnerella is used throughout the manuscript. All vaginal, cervical and endometrial samples, including negative controls were subjected to qPCR. AVM was defined as the presence of above threshold loads of Gardnerella and/or F. vaginae by qPCR as described previously(13). In addition, the total bacterial load was estimated in all the paired samples in oocyte donors by universal 16S qPCR(32). As some plates needed to be re- analyzed in the IVF cohort, insufficient amounts of purified sample were available for total bacterial load determination.

## Statistics

Table 1 was made in Stata (version IC 16.1), StataCorp LLC. All other analyses were performed in R (version 4.0.2) as described below. Bacterial load was expressed as 16S rRNA gene copy numbers per sample. Table 2 was made using a two-part model in the lme4 package(33). Part-1 of the model included the zero values and dichotomized outcome into zero or non-zero. A generalized linear mixed effect model with binomial-family was then used with fixed effects according to sample site and random effect according to individual patient. This was done to calculate the probability of observing zeros in the given sample site. Part-2 of the model used the log- transformed non-zero values in a linear mixed effect model estimating the median number of 16S rRNA gene copies in the given sample site and its 95% confidence interval. This was because there were a lot of zero values particularly in the species-specific assays. For the comparison, the ratio between the medians of the respective sample sites is given. The numbers and ratios reported in table 2 are thus conditioned on being non-zero values. Non-parametric tests as indicated in legends of tables and figures were also used to compare qPCR counts between groups using either paired or non-paired tests, where applicable. For the non-parametric tests, the qPCR zero values were assigned the value 0.5 in all calculations and supplementary figures. When pairwise testing was performed across multiple groups, multiple testing correction was made, using the false discovery rate (FDR) method (Benjamini-Hochberg).

Heatmaps were constructed in the Complex heatmaps package v2.4.3(34). The ASV table was converted into heatmaps of individual patients with relative abundances of the top 20 ASVs in the entire dataset, **Figure 3 and 4**. The median proportion of reads in the top 20 ASVs was 85% IQR (58%-98%), range 8%-100% of sequence reads per sample. Hierarchical clustering was performed using the Jaccard distance metric and the hclust function with Ward.D2 linkage. Beta-diversity was assessed by principal component analysis (PCA) made on centered log-ratio transformed data and with 95% confidence interval ellipses in the mixOmics package, v6.12.2(35). We used the procrustes and protest functions in vegan(36) for permutational correlation test in the PCA of each sampling site. In order to investigate the variance between sample sites, we used the adonis function (vegan(36)) on both the Bray-Curtis (abundance of ASVs) and the Jaccard (presence/absence of ASVs) distance matrices for permutational multivariate analysis of variance (PERMANOVA). We adjusted for repeated measures per person using the strata argument. To investigate whether assumptions for using adonis were met, we first used the multivariate homogeneity of groups dispersion test betadisper with anova in R. In both procrustes and adonis 999 permutations were used. The ggplots2 package(37) was used to generate all figures, unless stated otherwise.

## Acknowledgements

We acknowledge the clinicians aiding in sampling from The Fertility Clinic, Skive Regional Hospital, Denmark and Instituto Bernabeu, Alicante, Spain and laboratory technicians at Statens Serum Institut for laboratory analyses, including purification, qPCR and sequencing. We thank Copan S.P.A., Brescia, Italy for providing the ESwab® and eNat® kits used in the study. Finally, we thank all participating patients.

## Contribution to authorship

TH, PH and JSJ designed the study. TH, PH, AB and BL recruited the patients. TH drafted the manuscript. AI, EP, TH and JSJ made the bioinformatic work-up. All authors read, revised and accepted the final manuscript for publication.

## Details of ethical approval

This study was approved by the Scientific Ethical committee, Central Denmark Region (M-2015- 345-15) and Scientific Ethical committee, Instituto Bernabeu, Alicante, Spain (.20/019Tut). Moreover, we prospectively registered both cohorts in clinicaltrials.gov, NCT03363828 and NCT04393857.

## Data availability statement

The 16S rRNA gene sequencing data will be deposited at the Sequence Read Archive (SRA) upon acceptance. All other data, supporting the findings of this study, including clinical data, are available from the corresponding author upon request: participant-level personally identifiable data are protected under the Danish Data Protection Act and European Regulation 2016/679 of the European Parliament and of the Council (GDPR) which prohibit distribution even in pseudo- anonymized form, but can be made available under a data transfer agreement as part of a collaboration effort.

